# Machine Learning Prediction of Incidence of Alzheimer’s Disease Using Large-Scale Administrative Health Data

**DOI:** 10.1101/625582

**Authors:** Ji Hwan Park, Han Eol Cho, Jong Hun Kim, Melanie Wall, Yaakov Stern, Hyunsun Lim, Shinjae Yoo, Hyoung Seop Kim, Jiook Cha

## Abstract

Nationwide population-based cohort provides a new opportunity to build a completely automated risk prediction model based on individuals’ history of health and healthcare beyond existing risk prediction models. We tested the possibility of machine learning models to predict future incidence of Alzheimer’s disease (AD) using large-scale administrative health data. From the Korean National Health Insurance Service database between 2002 and 2010, we obtained de-identified health data in elders above 65 years (N=40,736) containing 4,894 unique clinical features including ICD-10 codes, medication codes, laboratory values, history of personal and family illness, and socio-demographics. To define incident AD two operational definitions were considered: “definite AD” with diagnostic codes and dementia medication (n=614) and “probable AD” with only diagnosis (n=2,026). We trained and validated a random forest, support vector machine, and logistic regression to predict incident AD in 1,2,3, and 4 subsequent years. For predicting future incidence of AD in balanced samples (bootstrapping), the machine learning models showed reasonable performance in 1-year prediction with AUC of 0.775 and 0.759, based on “definite AD” and “probable AD” outcomes, respectively; in 2-year, 0.730 and 0.693; in 3-year, 0.677 and 0.644; in 4-year, 0.725 and 0.683. The results were similar when the entire (unbalanced) samples were used. Important clinical features selected in logistic regression included hemoglobin level, age, and urine protein level. This study may shed a light on the utility of the data-driven machine learning model based on large-scale administrative health data in AD risk prediction, which may enable better selection of individuals at risk for AD in clinical trials or early detection in clinical settings.

## Introduction

Screening individuals at risk for Alzheimer’s disease (AD) based on medical health records in preclinical stages may lead to early detection of AD pathology and to better therapeutic strategies for delaying the onset of AD.^1–3^ Current biomarkers of AD requires the collection of specimen (e.g., serum or fluid) or imaging data. On the other hand, the electronic healthcare data, such as health records in clinical settings, or administrative health data, does not require additional time or effort for data collection. Also, with the advent of digitalization the amounts of such data have exponentially increased.^4^ Since it is ubiquitous, cost-effective and enormous, the digitalized healthcare database may be an invaluable resource for testing scalable predictive models for AD and other diseases alike. However, despite of its tremendous potential value, little is known about the extents to which the large-scale administrative health data is useful in AD risk prediction.

For AD risk prediction, prior models are typically based on predefined health profile variables, such as sociodemographic (age, sex, education), lifestyle (physical activity), midlife health risk factors (systolic blood pressure, BMI and total cholesterol level);^5,6^ and cognitive profiles.^7,8^ An important outstanding question is whether those simple predictive models based on the small sets of selected variables may sufficiently account for the heterogeneous etiologies of multi-factorial AD in clinical settings. Indeed, a meta-analysis study shows that multi-factor models best predict risk for dementia, whereas single-factor models do poorly,^6^ suggesting accurate AD risk prediction requires a large feature space. Here we test the extents to which a data-driven machine approach harvests salient information from the large-scale healthcare data containing thousands of data of individuals’ health trajectories and make an individual-specific prediction of AD risk.

Machine learning is an optimal choice of analytics for analyzing the large-scale administrative health data containing thousands of descriptors from hundreds of thousands of individuals. Studies show successful applications of machine learning to the nationwide administrative data in predicting incident diseases other than AD (diabetes, metabolic syndrome, suicide death, opioid overdose or drug-resistant epilepsy, etc).^9–13^ Given the recent rapid growth of the machine learning technology, application of the AI technology to clinical predictive modeling is likely to have a deep impact on medicine.^14–16^ But to our knowledge the data-driven predictive modeling based on nationwide population-based administrative health data has yet to be tested in AD risk prediction.

In testing predictive models, it is important to use sufficiently large data representative of the population. The data is important for the model accuracy, while the representativeness is important for the model generalizability. In this study, we used the National Health Insurance Service–national sample cohort (NHIS-NSC) of one million people representative of the South Korean population within the Korean National Health Insurance Service database^17^. Using the large-scale, thorough, longitudinal, administrative healthcare data (e.g., insurance claims and health check-ups) within this database, we constructed and validated data-driven machine learning models to predict future incidence of AD.

## Materials and Methods

### Datasets

NHIS-NSC cohort consist of randomly selected 1,025,340 participants comprising 2.2% of the total eligible Korean population in 2002, and followed for 11 years until 2013 unless participants’ eligibility was disqualified due to death or emigration^17^. This database contains for each individual’s features of services, diagnoses, and prescriptions associated with all the health care services provided by the NHIS. Clinical features include demographics and income levels divided by 10 levels based on subject’s monthly salary from the *Participant Insurance Eligibility database*; disease and medication codes from the *Healthcare Utilization database*; and laboratory values, health profiles, and history of personal and family illness from the *National Health Screening database* (from bi-annual health check-up required for elders with age above 40). Of those samples, 40,736 elders were selected in this study, whose records exist in all the three databases (Participant Insurance Eligibility database, Healthcare Utilization database, and National Health Screening database).

### Operational Definition of AD

For an operational definition of AD, a study of Canadian EMR from 3,404 adults shows sensitivity of 79% and specificity of 99% when they used an algorithm of “one hospitalization code OR three physician claims codes at least 30 days apart in a two year period OR a prescription filled for an AD-RD specific medication”^18^. In this study, to further improve the accuracy of an operational definition of AD, particularly sensitivity, we used the following algorithm to operationally define incident AD, herein labelled as *definite AD:* ICD-10 codes of AD^19^ (F00, F00.0, F00.1, F00.2, F00.9, G30, G30.0, G30.1, G30.8, G30.9) AND dementia medication prescribed with an AD diagnosis (e.g., donepezil, rivastigmine, galantamine, and memantine). Furthermore, we considered a broader definition of AD using only ICD-10 codes to minimize false negative cases (e.g. individuals with AD diagnose who did not take medication); this was labeled as *probable AD.* Within each individual with either definition of incident AD, the data after the incidence was excluded. Based on these two operational definitions, the prevalence rates were 1.5% for definite AD and 4.9% for probable AD; the former was smaller than what is reported in a door-to-door visit study in Korean elders (age > 65 years old), but the latter was similar to that^20^.

### Data and Preprocessing

We used the following variables from the NHIS-NSC data: 21 features including laboratory values, health profiles, history of family illness from the Health Screening database; 2 features including age and sex from the Participant Insurance Eligibility database; and 6,412 features including ICD-10 codes and medication codes. Descriptions of data coding and exclusion criteria for all the features except for ICD-10 codes and medication codes are available in **Supplementary Table 1.**

Our data preprocessing steps are as follows. (i) Data alignment: We aligned the data to each individual’s initial AD diagnosis (event-centric ordering). (ii) ICD-10 and medication coding: Since ICD-10 and medication codes have hierarchical structures, we used the first disease category codes (e.g., F00 [Dementia in Alzheimer’s disease] including F00.0 [Dementia in Alzheimer’s disease with early onset], F00.1 [Dementia in Alzheimer’s disease with late onset], F00.2 [Dementia in Alzheimer’s disease, atypical or mixed type], and F00.9 [Dementia in Alzheimer’s disease, unspecified]), and the first 4 characters for the medication codes representing main ingredients. (iii) Rare disease or medication codes found less than five times in the entire data were excluded from the analysis (1,179 disease and 362 medication codes). (iv) if a participant has no health screening data (laboratory values, health profiles, and history of personal and family illness from the National Health Screening database) during the last two years of the processed data (in Korea a biannual health screening is required for every elder), we excluded that participant from the analysis. This preprocessing procedure yielded 4,894 unique variables used in the models (see **Supplementary Table 2** for detailed information).

For each *n*-year prediction, within the AD group, we used the data between 2002 and the year of incident AD – *n* because it requires at least *n* years prior to the incident AD. Within the non-AD group, we used the data from 2002 to 2010 – *n*. For example, for 0 year prediction, if a patient was diagnosed with AD at 2009, we used the data between 2002 and 2009; for 1 year 2002-2008; for 2 year prediction, 2002-2007; for 3 year, 2002-2006; and for 4 year, 2002-2005.

For model training, validation, and testing, we used the randomly sampled balanced dataset, as well as the entire, unbalanced dataset. For the balanced dataset, we performed bootstrap sampling with replacement 10 times.

### Machine learning analysis

We implemented three machine learning algorithms: random forest, support vector machine with linear kernel, and logistic regression. Model training, validation, and testing was done using nested stratified 5-fold cross validation with 5 iterations. Feature selection was done within train sets using the variance threshold method.^21^ Hyper-parameters optimization was done within validation sets. The following hyper-parameters were tuned: for random forest, the minimum number of samples required at a leaf node and the number of trees in the forest; for support vector machine, regularization strength; for logistic regression, the inverse of regularization strength. In logistic regression L2 regularization was used. Lastly, generalizability of model performance was assessed on the test sets. We measured the following model performance metrics in the test set: The area under the receiver operating characteristic curve (ROC), sensitivity and specificity.

### Data availability

Codes are available at https://github.com/a011095/koreanEHR. The data in this study is available upon request.

### Ethical approval

This study complies with the Transparent Reporting of a Multivariable Prediction Model for Individual Prognosis or Diagnosis (TRIPOD) reporting guideline. The study with exemption of informed consent (for retrospective, de-identified, publicly available data) was approved by the Institutional Review Board of National Health Insurance Service (NHIS) Ilsan Hospital, Gyeonggi-do, Korea (IRB number NHIMC 2018-12-006). All methods in this study were performed in accordance with the Declaration of Helsinki.

## Results

### Sample characteristics

Of 40,736 individuals with age above 65 years in 2002, we identified 614 unique individuals with AD incidence using the definite AD outcome, 2,026 with AD incidence using the probable AD definition, and 38,710 elders with no AD incidence. The rate of AD in this cohort was 1.56% using the definite AD definition, and 4.97% using the probable AD definition. Demographic characteristics showed significant differences in age between both AD groups and non-AD groups and non-significant differences in income and sex (**Table 1**).

**Table 1.**
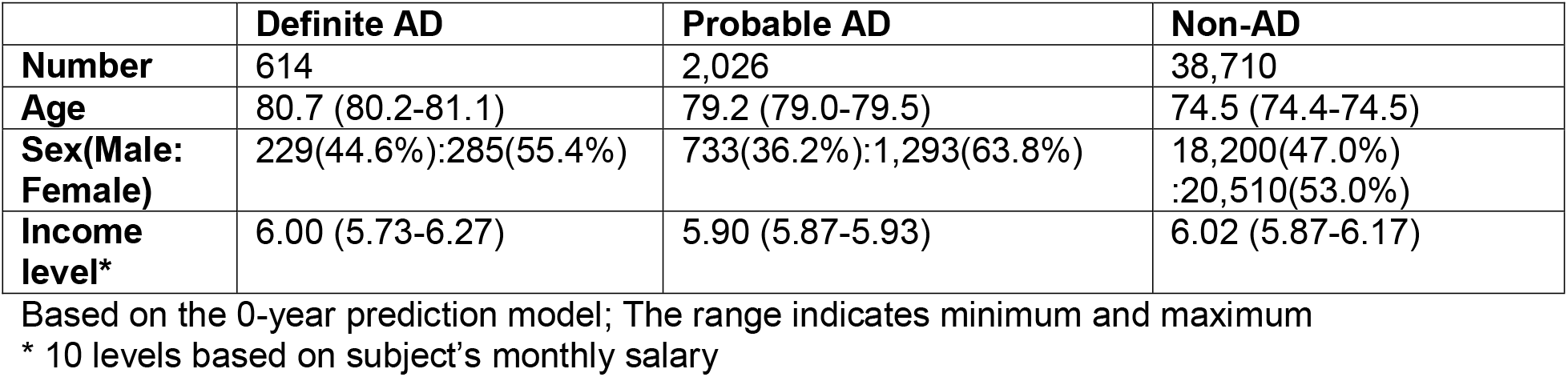
Sample characteristics

|  | **Definite AD** | **Probable AD** | **Non-AD** |
| --- | --- | --- | --- |
| **Number** | 614 | 2,026 | 38,710 |
| **Age** | 80.7 (80.2-81.1) | 79.2 (79.0-79.5) | 74.5 (74.4-74.5) |
| **Sex(Male:**<br>**Female)** | 229(44.6%):285(55.4%) | 733(36.2%):1,293(63.8%) | 18,200(47.0%)<br>:20,510(53.0%) |
| **Income**<br>**level\*** | 6.00 (5.73-6.27) | 5.90 (5.87-5.93) | 6.02 (5.87-6.17) |

Based on the 0-year prediction model; The range indicates minimum and maximum
\* 10 levels based on subject's monthly salary

### Model prediction

Classifiers were trained to predict 0,1,2,3, and 4 subsequent-year incident AD. In balanced samples (bootstrapping with replacement) (**Table 2**), when using the definite AD definition (based on ICD-10 codes and dementia prescription), in predicting 0yr incidence of AD, random forest (RF) showed the best performance with accuracy 0.823 and AUC of 0.898. When using the probable AD definition (based on ICD-10 codes), classification performance was slightly lower with accuracy of 0.788 and AUC of 0.850 (RF). Classification performance decreased in predicting future incident AD of later years: using the definite AD definition, accuracy/AUC of 0.713/0.775(1 year), 0.675/0.730(2 year), 0.632/0.677(3 year), 0.663/0.725(4 year); using the probable AD definition, accuracy/AUC of 0.688/0.759(1 year), 0.645/0.693(2 year), 0.610/0.644(3 year), 0.641/0.683(4 year). The results were similar when we used the entire, unbalanced samples for model training and evaluation **(Supplementary table 3)**, RF showed the best performance in predicting 0yr incidence of AD with AUC of 0.887 when using the definite AD definition and AUC of 0.805 when using the probable AD definition. Classification performance decreased as the predicting period getting longer; using the definite AD definition, AUC of 0.781 (1 year), 0.739 (2 year), 0.686 (3 year), and 0.662 (4 year); using the probable AD definition, AUC of 0.730 (1 year), 0.645 (2 year), 0.575 (3 year), and 0.602 (4 year). Numbers of features and look-back periods also decreased in later year (**Supplementary Table 2**).

**Table 2.** Performance of AD predictive models trained on NHIS-NSC by using balanced samples

| Definite AD ( AD/ non-AD 614/614) |  |  |  |  |  |
| --- | --- | --- | --- | --- | --- |
|  | Classifier | Accuracy | AUC | Sensitivity | Specificity |
| 0 yr | LR | 0.76 | 0.794 | 0.726 | 0.793 |
|  | SVM | 0.763 | 0.817 | 0.715 | 0.811 |
|  | RF | 0.823 | 0.898* | 0.795 | 0.852 |
| 1 yr | LR | 0.677 | 0.693 | 0.65 | 0.704 |
|  | SVM | 0.678 | 0.705 | 0.699 | 0.656 |
|  | RF | 0.713 | 0.775* | 0.686 | 0.74 |
| 2 yr | LR | 0.652 | 0.684 | 0.639 | 0.666 |
|  | SVM | 0.663 | 0.687 | 0.572 | 0.753 |
|  | RF | 0.675 | 0.730* | 0.608 | 0.742 |
| 3 yr | LR | 0.623 | 0.645 | 0.562 | 0.684 |
|  | SVM | 0.607 | 0.635 | 0.58 | 0.633 |
|  | RF | 0.632 | 0.677* | 0.572 | 0.693 |
| 4 yr | LR | 0.627 | 0.661 | 0.509 | 0.745 |
|  | SVM | 0.646 | 0.685 | 0.538 | 0.754 |
|  | RF | 0.663 | 0.725* | 0.621 | 0.705 |
| Probable AD ( AD/ non-AD 2026/2026) |  |  |  |  |  |
|  | Classifier | Accuracy | AUC | Sensitivity | Specificity |
| 0 yr | LR | 0.736 | 0.783 | 0.689 | 0.783 |
|  | SVM | 0.734 | 0.794 | 0.652 | 0.816 |
|  | RF | 0.788 | 0.850* | 0.723 | 0.853 |
| 1 yr | LR | 0.663 | 0.697 | 0.634 | 0.692 |
|  | SVM | 0.661 | 0.691 | 0.592 | 0.729 |
|  | RF | 0.688 | 0.759* | 0.609 | 0.767 |
| 2 yr | LR | 0.643 | 0.672 | 0.633 | 0.654 |
|  | SVM | 0.645 | 0.68 | 0.58 | 0.709 |
|  | RF | 0.638 | 0.693* | 0.564 | 0.713 |
| 3 yr | LR | 0.61 | 0.635 | 0.557 | 0.663 |
|  | SVM | 0.597 | 0.644* | 0.427 | 0.767 |
|  | RF | 0.581 | 0.609 | 0.505 | 0.657 |
| 4 yr | LR | 0.611 | 0.644 | 0.516 | 0.707 |
|  | SVM | 0.601 | 0.641 | 0.465 | 0.738 |
|  | RF | 0.641 | 0.683* | 0.603 | 0.679 |
| * best performing models based on AUC. |  |  |  |  |  |
| AD, Alzheimer's dementia ; LR, logistic regression; SVM, support vector machine; RF, random forest |  |  |  |  |  |

### Important features

Logistic regression identified the features positively related to incident AD. These included age (b = 0.689; odd ratio(OR) = 1.991), elevated urine protein (b = 0.303; OR = 1.353), prescription of Zotepine (antipsychotic drug) (b = 0.303; OR = 1.353), and the features negatively related to incident AD, such as, decreased hemoglobin (b = −0.902; OR = 0.405), prescription of Nicametate Citrate (b = −0.297; OR = 0.743), diagnosis of other degenerative disorders of nervous systems (b = −0.292; OR = 0.747), and disorders of the external ear (b = −0.274; OR = 0.760) (**Table 3**).

**Table 3.** Top ten features and weights from logistic regression (0-yr prediction).

| Type of data | Name | b value | 95% CI | Odd ratio | p-value |
| --- | --- | --- | --- | --- | --- |
| health checkup | Hemoglobin(g/dL) | -0.902 | -0.903/-0.901 | 0.405 | <0.001 |
| demography | age | 0.689 | 0.687/0.690 | 1.991 | <0.001 |
| health checkup | urine protein* | 0.303 | 0.300/0.306 | 1.353 | <0.001 |
| medication | Zotepine (antipsychotic drug) | 0.303 | 0.280/0.325 | 1.353 | <0.001 |
| medication | Nicametate Citrate (vasodilator) | -0.297 | -0.298/-0.295 | 0.743 | <0.001 |
| disease code | other degenerative disorders of nervous system in diseases classified elsewhere | -0.292 | -0.309/-0.274 | 0.746 | <0.001 |
| disease code | disorders of external ear in diseases classified elsewhere | -0.274 | -0.328/-0.220 | 0.760 | <0.001 |
| medication | Tolfenamic acid 200mg (pain killer) | -0.266 | -0.279/-0.254 | 0.766 | <0.001 |
| disease code | adult respiratory distress syndrome | -0.259 | -0.282/-0.236 | 0.771 | <0.001 |
| medication | Eperisone Hydrochloride (antispasmodic drug) | 0.255 | 0.237/0.272 | 1.290 | <0.001 |

\*Urine protein was detected by urine dipstick test (1 : negative (-), 2 : weak positive (±), 3 : positive (1+), 4 : positive (2+), 5 : positive (3+), 6 : positive (4+))

### Model prediction using important features only (Table 2)

After identified the important features related to incident AD by logistic regression, classifiers were trained with top 20 important features only to predict 0,1,2,3 and 4 subsequent-year incidence of AD. The AUC were higher when using all features than when using top 20 features. In longer periods, the opposite result was obtained.

## Discussion

This study assessed the utility of the nationwide population-based administrative health data in predicting the future incidence of AD. Using machine learning, we predicted future incidence of AD with acceptable accuracy of 0.713 (in terms of AUC 0.781) in one-year prediction. The high accuracy of our models based on large nationwide samples may lend a support to the potential utility of the administrative data-based predictive model in AD. Despite of the limitations inherent to the administrative health data, such as the inability to directly ascertain clinical phenotypes, this study demonstrates its potential utility in AD risk prediction, when combined with data-driven machine learning.

Our model performance with AUC of 0.898, 0.775, and 0.725 in predicting baseline, subsequent one-year, and four-year incident AD is relatively accurate compared with the literature. In all-cause dementia risk prediction based on genetic (ApoE) or neuropsychological evaluations, MRI, health indices (diabetes, hypertension, lifestyle), and demographic (age, sex, education) variables, prior models show accuracy ranging from 0.5 to 0.78 in AUC (reviewed in ^22^). Of note, compared with these studies, our approach is solely based on administrative health data without neuropsychological, genetic testing, or brain imaging. This has important implications for the practical utility of the administrative dat -based risk prediction, in that it can provide an early indication of AD risk to clinicians. Together with existing screening tools (e.g., MMSE), this may assist deciding when to seek a further clinical assessment to a given patient in an individual-specific manner.

Comparing the models based on the sampled, balanced set and on the entire, unbalanced set shows a small-to-moderate difference in model performance. For example, based on the RF model in predicting 0-yr definite AD, the AUC’s are 0.887 and 0.898 in the unbalanced and balanced samples, respectively, showing a 1% increase. On the other hand, in predicting 4-yr definite AD, the AUC’s are 0.662 and 0.725 in the unbalanced and balanced samples, respectively, showing a 9.5% increase. These results show a trivial-to-moderate difference in model performance between balanced and balanced samples. However, we should point that, if one uses an algorithm capable of processing the temporal information among the clinical features, such as recurrent neural networks^23^, then using the entire data for scalable learning is likely to be beneficial.

Comparing the model performance across years, the 3-year prediction is less accurate than the 4-year prediction. This seems counter-intuitive at first, but our data shows that the length of data is greater in 4-year prediction than in 3-year prediction (**Supplementary Table 2**). We thus suspect that this difference in data availability may be a cause of the expected performance increase in later year prediction. This might be also related to the irregularity of the NHIS-NSC dataset due to changes in healthcare policy.

Our model detected interesting clinical features associated with incident AD. The data-driven selection of features is consistent with risk factors found in the literature. A decrease in hemoglobin level was selected as the feature most strongly associated with incident AD. Indeed, anemia is known as an important risk factor for dementia.^24–26^ A study using National Health Insurance Service-National Health Screening Cohort (NHIS-HEALS), the NHIS health screening data in Korea, not only found that anemia was associated with dementia, but also revealed a dose-dependent relationship between anemia and dementia.^27^ Likewise, our data-driven model shows the hemoglobin level as the most significant predictor. This finding has implications for public health because anemia is a modifiable factor. Given our finding and the consistent literature on the large association between hemoglobin level and AD and other dementia, future research may investigate the biological pathway of anemia’s contribution to AD pathology and cognitive decline.

We also noted a positive association between urine protein level and incident AD. In the NHIS-NSC, protein in urine is typically measured using urine dip stick. This approach is not a quantitative measure of urine protein, but it is useful as a screening method for proteinuria.^28,29^ Literature shows association between albuminuria and dementia.^30^ Our finding suggests the potential utility of a urine test as part of the routine health check-up in AD risk prediction.

Four medications were also associated with incident dementia within top ten features. We found that Zotepine, Eperisone hydrochloride had a positive association and Nicametate Citrate and Tolfenamic acid had a negative association with incident AD. It is interesting that patients prescribed tolfenamic acid showed lower incidence of AD.

This drug used in Korea for pain control in conditioner such as rheumatoid arthritis. It is known to lower the gene expression of Amyloid precursor protein 1(APP1) and beta-site APP cleaving enzyme 1(BACE1) by promoting the degradation of specificity protein 1(Sp1).^31–33^ As a potential modifier of tau protein, Tolfenamic acid is under investigation as a potential drug to prevent and modify the progression of AD.^34^ The results of this study support the above experimental result and show that tolfenamic acid may be a potential anti-dementia medication.

Zotepine is an atypical antipsychotic drug with proven efficacy for treatment of schizophrenia. Our model showed the use of zotepine positively correlated with incident AD. There are two possible interpretations. Zotepine may have been used to treat behavioral and psychological symptoms of dementia (BPSD) before incident AD or diagnosis of AD.^35^ Thus, the prescription of Zotepine may indicate early AD symptoms and, consequently, an increasing likelihood of incident AD. Alternatively, some studies indicate that individuals with schizophrenia may have an increased risk for the development of dementia.^36^ Given this, it might be possible that the incident AD is high in individuals with schizophrenic symptoms to whom Zotepine is prescribed. However, this alternative interpretation may be questionable considering that, in our model, the disease code of Schizophrenia has not been selected as an important feature. In either case, it should be noted that, while our results indicate a potential relationship between Zotepin and incident AD (likely reflecting the common practice in dementia), no causal relationship should be drawn.

Nicametate Citrate, a vasodilator, was also negatively associated with incident AD. This may be in line with the literature showing effects of vasodilators on increasing cognitive function and reducing the risk of vascular dementia, although the exact mechanism remains unclear.^37,38^ Further research is required.

## Limitations

One of the limitations of this study is that diagnoses of AD in our database are not clinically ascertained. For example, there may be incorrect diagnoses or misdiagnoses of AD in the claim data. To mitigate this issue, we firstly confirmed the similar prediction results using two different definitions of incident AD, “probable AD” (based on AD disease codes) and “definite AD” (based on both AD disease codes and anti-dementia medication). Secondly, in South Korea, every elder with age 60 years old is required to have complementary dementia screening supported by the National Health Insurance Service at public healthcare centers, where individuals that high-risk for dementia get referred to physicians for further clinical examination. Such a system may help reduce false negative cases. Lastly, Korean health insurance system and policies support the reliability of the AD diagnoses. That is, the Health Insurance Review and Assessment Service of NHIS reviews and supervises the medical claims of AD medication^i^ Thus, it is likely that individuals with records of receiving dementia medication meet strong diagnostic criteria. These aspects may alleviate potential validity issues of the AD diagnoses in the Korean administrative health data. Another limitation is that the features associated with incident AD do not indicate causality. Rather, this finding indicates a data-driven discovery from a machine learning model trained on large administrative data, which might be useful to generate new hypotheses, to confirm existing ones, or to compare relative importance in predicting incident AD considering large feature space. We believe this is a useful value of data-driven science.

## Conclusions

In sum, this study lends support to a statistically meaningful detection of individuals with AD risk solely based on the administrative health data. Generalizability of our findings to independent data in other nations, ethnicities, and healthcare and insurance systems remains to be tested. If replicated, this study may further motivate the implementation of a system in clinical settings that could alarm a risk for AD, which may enable earlier and more accurate screening for subsequent clinical testing.

**Figure 1.**
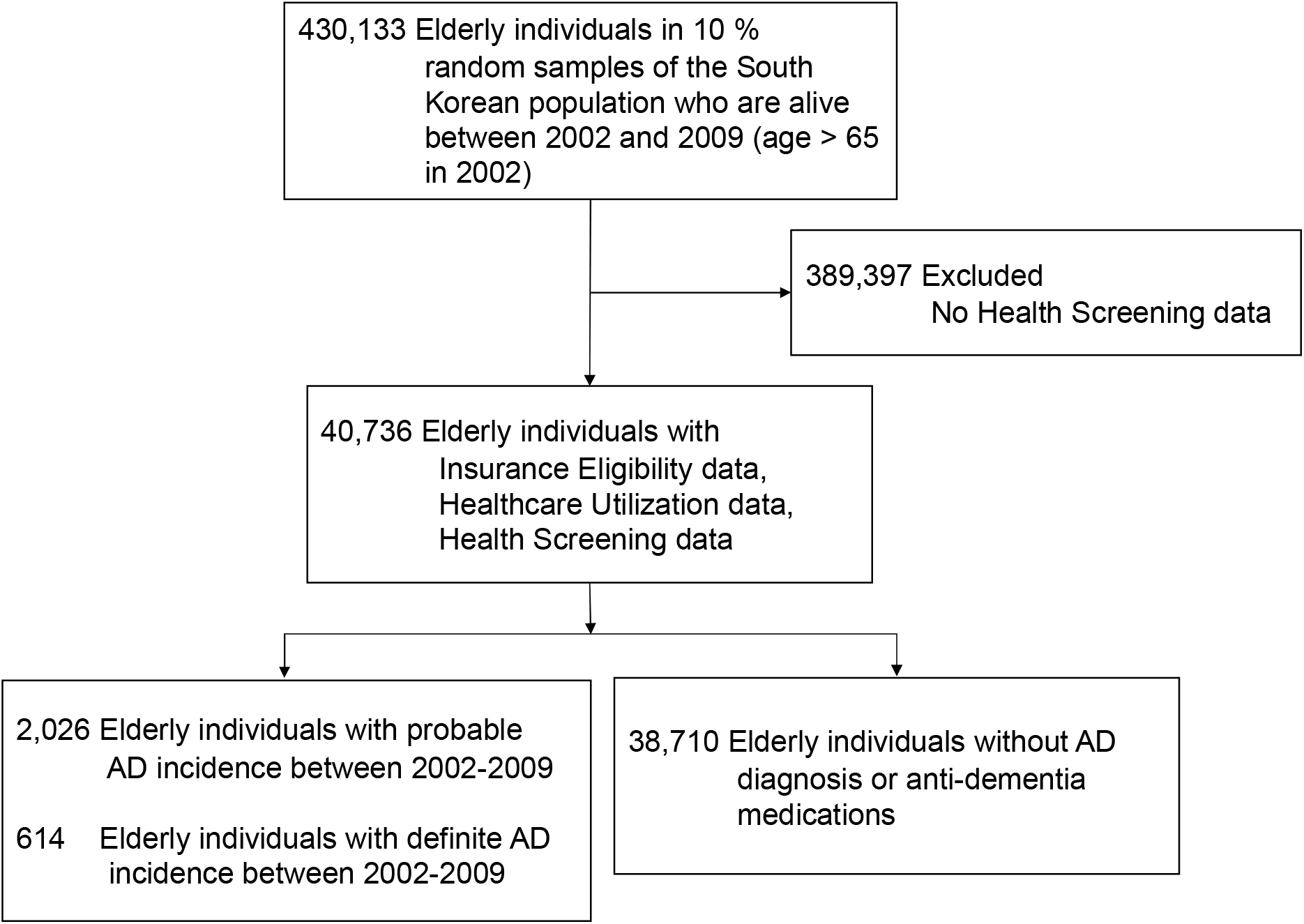
Consort Diagram.

**Figure 2.**
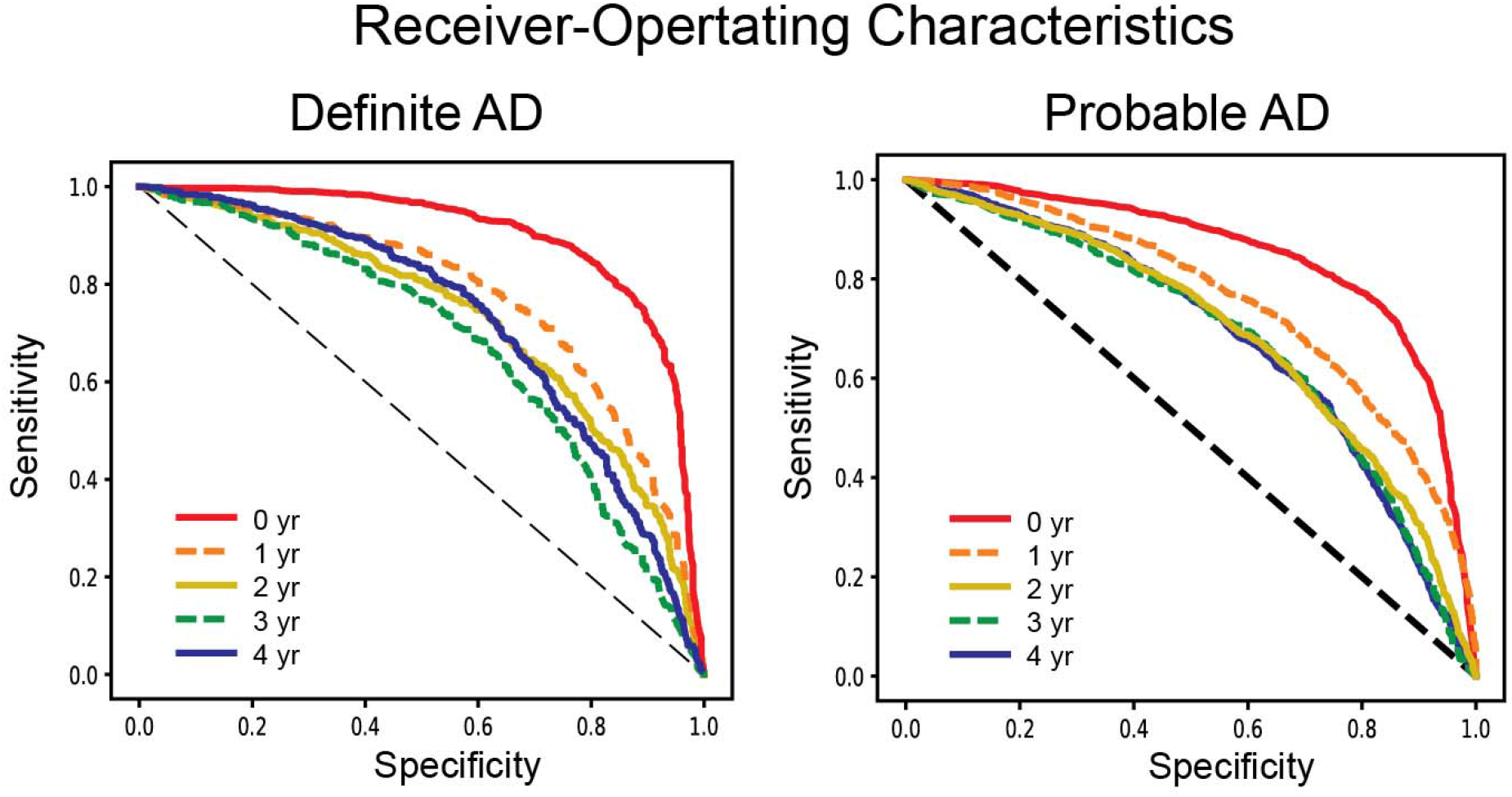
Performance of machine learning models in predicting incident AD. Receiver-Operating Characteristic plots are shown for 0,1,2,3,4-subsequent year prediction. Incident AD was defined based on ICD-10 AD codes and anti-dementia medication for AD, “Definite AD”, or based on AD codes only, “Probable AD”. In each year prediction, a best performing model was selected for plotting.

## Acknowledgement

This study is funded by NHIS Ilsan Hospital Research Support Program (PI: Kim, HS); National Institute of Mental Health through K01-MH109836 (PI: Cha); Brain Behavior Research Foundation Young Investigator Award (PI: Cha); Korean Scientists and Engineers Association Young Investigator Grant (PI: Cha); Brain Pool Program through the National Research Foundation of Korea (NRF) funded by the Ministry of Science and ICT (200-20190251; PI: Cha).

## Author Contributions

Conception and design: H.S.K, J.C

Data collection and analysis: J.H.P, H.L, S.Y, J.H.K, H.E.C, H.S.K, J.C

Data interpretation: J.H.P, M.W, Y.S, S.Y, H.E.C, H.S.K, J.C,

Manuscript writing: H.E.C, J.H.P, H.S.K, J.C,

Revision after critical review: all authors

## Footnotes

i For example, it requires the following conditions to consider the insurance coverage of dementia medication: for donepezil and rivastigmine patches, MMSE (Mini-Mental State Examination) =< 26 and CDR (Clinical Dementia Rating) = 1~3 or GDS (Global Deterioration Scale)= 3~7; for galantamine and rivastigmine capsules, MMSE = 10 ~ 26 and CDR = 1~2 or GDS = 3~5; for memantine, MMSE =< 20 and CDR = 2~3 or GDS = 4~7 (**Supplementary Figure 1**).

